# EditorForge: An Active-Site-Aware Framework for Inverse-Folding-Based Protein Redesign

**DOI:** 10.64898/2026.06.08.730993

**Authors:** Anthony Chen, Jibraan Siddiqui, William Taucar, Leo Tiralongo, Maxim Tkachenko, Andy Xu, Sehaj Bawa, Sihan Guo, Owen Pinska, Jonathan Rim, Joshua Shi, Michael Wang, Eason Zhao

## Abstract

Inverse-folding models can rapidly generate protein sequences compatible with a supplied backbone, but unconstrained redesign is poorly suited to enzyme and genome-editor-associated domains, where catalytic, substrate-proximal, and conserved structural regions must remain protected. In this paper, we present EditorForge, a modular constraint-and-audit suite for editor-domain protein redesign that wraps fixed-backbone inverse folding with explicit design masks, fixed-position enforcement, active-site-proximity auditing, active-site-shielded regeneration, and downstream structural quality control. Using full-length Moloney murine leukemia virus reverse transcriptase structure 4MH8 (MMLV RT 4MH8) as a demonstration target, EditorForge first restricted redesign to a bounded 25-position envelope while fixing 428 residues. An initial audit detected active-site-proximal failure modes despite fixed-position integrity. Later, the Active Site Shield module then removed five unsafe design positions, replaced them with lower-contact alternatives, and regenerated candidates under stricter constraints. Post Shield Audit evaluated 24 regenerated candidates, all of which satisfied the hard sequence/mask and active-site-shield constraints. For the eight candidates that were selected or returned for structure-prediction/refolding quality control, Enhanced RefoldQC found that all 8 evaluated predicted structures passed the computational structure-QC screen. That said, the selected 8 candidates passed the computational structure-QC screen, with global C*α* RMSD values of 1.2061–1.5555 °A, active-site C*α* RMSD values of 0.4098–1.8397 °A, mutation-neighborhood C*α* RMSD values of 1.3155-1.6848 °A, and average pLDDT-like confidence values of 94.87–95.11. In short, EditorForge provides a reproducible triage layer that converts general inverse-folding output into constrained and editor-specific candidate sets for downstream structural and biological review on top of existing structural prediction tools.

## 1 Introduction

Inverse-folding models have made fixed-backbone protein design much easier to start than before. [1, 2, 3, 4, 6, 8]. Given a protein structure, models such as ProteinMPNN can propose sequences predicted to be compatible with that backbone which is incredibly useful when the goal is to redesign selected positions within an existing protein domain. In that setting, structure-prediction tools such as AlphaFold, ColabFold, and ESMFold can provide a second layer of computational review by asking whether designed sequences are still predicted to fold into structures close to the parent model in the first place [9, 10, 11, 12].

For enzyme and genome-editor-associated domains, this general workflow is incomplete because structural compatibility is only one part of the design problem. Reverse transcriptases, polymerases, nucleases, deaminases, and editor-fusion components also contain catalytic residues, substrate-proximal regions, conserved motifs, nucleic-acid-interaction surfaces, and local structural features that may be essential for function [23, 22]. Because of this, a sequence can look acceptable to an inverse-folding model and still be a poor engineering candidate if it changes residues near the active site or disrupts positions that should have remained fixed, which is a huge practical problem because broad sequence redesign can produce candidates that are computationally plausible but biologically difficult to justify and thus difficult to be applicable to real world problems.

In particular, this gap matters especially for editor-domain protein engineering as in many redesign problems, the central question is whether the sequence was generated inside a biologically meaningful design envelope on top of the fact that if it can fit inside an existing backbone [14, 15]. As such, questions such as: which residues were allowed to change, which residues were forced to remain fixed, did any candidate mutate outside the permitted region, did any apparently acceptable candidate place mutations too close to catalytic-reference residues, all arise and without answering these questions explicitly, inverse-folding output can be difficult to interpret, even when the generated structures appear reasonable on paper Because of this, EditorForge was developed specifically for this intermediate layer, a constraint-and-audit suite for editor-domain redesign that wraps fixed-backbone inverse folding with a declared design mask, fixed-position enforcement, mutation auditing, active-site-proximity checks, active-site-shielded regeneration, and downstream structure-quality control with the goal to turn inverse folding from broad sequence generation into a controlled triage workflow for sensitive enzyme-like domains [6, 7].

In the EditorForge workflow, the parent sequence and parent structure are first converted into a design mask then the mask separates residues into designable and fixed positions then the designable set defines the allowed search region, while the fixed set protects the rest of the domain. Afterwards, candidate sequences are then generated under fixed-position constraints using ProteinMPNN. After generation, each candidate is compared against the parent sequence and candidates are then flagged or rejected if they mutate fixed residues, mutate outside the declared design envelope, or otherwise violate the intended redesign logic.

A useful feature of this workflow is that failure is treated as information rather than as a terminal result as in an editor-domain setting, a candidate set may satisfy fixed-position integrity but still place mutations too close to catalytic-reference residues. EditorForge handles this through an Active Site Shield module which removes design positions that fall within a hard active-site distance shell or local sequence-neighborhood window, replaces them with lower-contact alternatives (sourced through online datasets and knowledge) outside the protected region, and reruns the design step under the stricter shielded mask. In this way, the workflow can detect a failure mode, update the constraint set, and regenerate candidates instead of accepting the first plausible-looking output, which has proven to be quite successful as shown further in this preprint.

We demonstrate EditorForge on Moloney murine leukemia virus reverse transcriptase using the full-length reverse-transcriptase structure 4MH8 as the structural target, a useful demonstration case because reverse transcriptase is structurally characterized, active-site-sensitive, and relevant to editor-associated protein engineering [16, 17]. In the demonstration workflow, EditorForge restricted redesign to a 25-position design envelope while fixing 428 residues. An initial constrained design pass preserved fixed-position integrity but still produced active-site-proximal risk, to which Active Site Shield then removed five unsafe design positions, replaced them with five lower-contact alternatives, and regenerated candidates under the stricter active-site-shielded mask.

The final shielded workflow produced 24 regenerated candidates, from which 8 were selected for structure-prediction and refolding quality control. Enhanced RefoldQC found that all 8 candidates passed the computational structure-QC screen and across the final candidates, global C*α* RMSD ranged from 1.2061 to 1.5555 °A, active-site C*α* RMSD ranged from 0.4098 to 1.8397 °A, mutation-neighborhood C*α* RMSD ranged from 1.3155 to 1.6848 °A, and average pLDDT-like confidence values ranged from 94.87 to 95.11. These results support the result that EditorForge can convert general inverse-folding output into constrained, auditable, editor-domain-specific candidate sets that preserve parent-like global and local geometry under computational refolding QC.

The contribution of this work is methodological as we do not claim that any generated candidate improves reverse-transcriptase activity, editing performance, fidelity, processivity, expression, delivery, or safety because all these require wet-lab proofing, none of which was conducted in this paper. Rather, EditorForge provides a reproducible computational layer between inverse-folding generation and downstream structural or biological review since it defines where redesign is allowed, detects when generated candidates stress or violate that design envelope, regenerates under corrected constraints, and reports candidates in a form suitable for further evaluation, whether it be further computaitonal pipelines or simply biological processes. Simulations, scripts, and further data can be found on GitHub technical repository: https://github.com/anthonycehn-eng/EditorForge-An-Active-Site-Aware-Framework-for-Inverse-Folding-Based-Protein-Redesign.git.

## 2 Methods

### 2.1 EditorForge workflow and constrained design setup

EditorForge was designed as a constraint-and-audit workflow for fixed-backbone redesign of editor-associated protein domains as well as a computational triage layer. The workflow starts from a parent amino-acid sequence and a parent structural model, defines a restricted set of residues that may be redesigned, generates candidate sequences under fixed-position constraints, audits the resulting candidates for violations of the declared design envelope, revises the design mask when active-site-proximal risk is detected, and forwards the surviving candidates to structure-prediction and refolding quality control.

The demonstration target was Moloney murine leukemia virus reverse transcriptase, using the full-length reverse-transcriptase structure 4MH8 as the parent structural model [16, 17, 18, 19]. The structure was converted into a single-chain working target with a corresponding structure-derived FASTA sequence. All residue positions reported in masks, mutation tables, active-site audits, candidate sheets, and RefoldQC outputs refer to this working coordinate system. The working parent domain contained 453 residues.

The constrained design setup partitioned the parent residue index set *P* = {1, …, *N*}, with *N* = 453, into a designable set *D* and a fixed set *F* = *P* \ *D*. Positions in *D* were allowed to vary during inverse-folding generation, whereas positions in *F* were expected to remain parent-identical. The final design envelope contained 25 designable residues and 428 fixed residues, so only a small fraction of the parent RT domain was made mutable.

The initial designable set was constructed using conservative structural and biochemical rules [6, 7]. For instance, terminal and domain-boundary regions were protected as well as chemistry-sensitive residue classes, including residues likely to be catalytic, metal-binding, conformationally special, or otherwise structurally sensitive, were also protected. For the remaining residues, EditorForge estimated local structural burial using C*α* contact counts. For residue *i*, the contact count was computed as

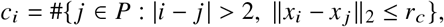

where *x*_*i*_ and *x* _*j*_ are C*α* coordinates and *r*_*c*_ is the contact-distance cutoff. Lower-contact residues were preferentially selected as candidate design positions, subject to spacing and protection constraints.

ProteinMPNN was used as the fixed-backbone inverse-folding engine under the fixed-position constraints exported by EditorForge [4]. Generated candidate sequences *s*^(*k*)^ were obtained through comparison against the parent sequence *s*^(0)^ and not accepted directly from ProteinMPNN score alone

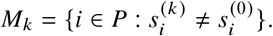

A candidate passed the primary integrity audit only if

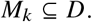

Any mutation in *F* was recorded as a fixed-position violation, and any mutation outside the declared design set was recorded as an outside-design mutation.

### 2.2 Active-site shielding, regeneration, and post-shield auditing

Fixed-position integrity alone is not sufficient for editor-domain redesign because a candidate can satisfy the declared design mask while still placing allowed mutations too close to catalytic or substrate-proximal regions. Therefore EditorForge performs an active-site-proximity audit after candidate generation. For the MMLV reverse-transcriptase demonstration, the active-site reference set was

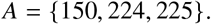

For each mutated or designable position *i*, EditorForge computed the nearest active-site-reference distance

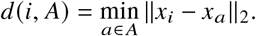

Positions within the hard active-site shield were considered unsafe for redesign. In the final workflow, the hard distance cutoff was 16 °A. A local sequence-neighborhood protection rule was also applied around the active-site reference residues using a ± 2 residue window.

When the initial constrained audit detected active-site-proximal design risk, EditorForge applied Active Site Shield. This module removed design positions that fell within the hard active-site distance shield or local active-site sequence window. Removed positions were replaced with lower-contact alternatives outside the hard shield so that the 25-position design budget was preserved. In the final MMLV RT workflow, Active Site Shield removed five unsafe design positions, added five replacement positions, and produced a revised 25-position shielded design mask.

ProteinMPNN was then rerun using the active-site-shielded mask, to which the regenerated candidates were evaluated by Post Shield Audit. This audit recorded mutation count, fixed-position violations, outside-design mutations, hard active-site-shield breaches, peripheral-zone mutations, nearest active-site mutation distance, ProteinMPNN score, shield score, and candidate tier. The post-shield audit evaluated 24 regenerated candidates and eight final predicted candidate structures were subsequently evaluated by Enhanced RefoldQC. These eight final candidates had zero fixed-position violations, zero outside-design mutations, and zero hard active-site-shield breaches in the final candidate dashboard.

### 2.3 Structure-prediction handoff, Enhanced RefoldQC, and reproducibility package

The final candidate set was exported as a structure-prediction packet containing a combined FASTA file, individual FASTA files, candidate metadata, and a manifest linking each sequence to its EditorForge candidate ID. The handoff was designed to be model-agnostic: predicted structures from ColabFold, ESMFold, AlphaFold-style workflows, or equivalent structure-prediction tools can be returned to the expected output directory and parsed by the same quality-control module [9, 10, 11, 12].

Enhanced RefoldQC evaluated whether predicted candidate structures preserved the parent fold and local geometry around sensitive regions. For each predicted PDB file, C*α* coordinates were extracted and aligned to the parent structure using Kabsch alignment [24, 25, 26, 27]. For coordinate sets *X* = (*x*_1_, …, *x*_*m*_)and *Y* = (*y*_1_, …, *y*_*m*_), aligned C*α* RMSD was computed as

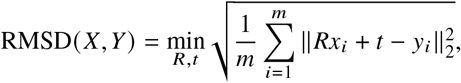

where *R* is a rotation matrix and *t* is a translation vector.

RefoldQC computed three structural metrics for each evaluated candidate: global C*α* RMSD measured full-chain similarity between the predicted candidate structure and the parent structure, active-site C*α* RMSD measured preservation of the catalytic-reference region, mutation-neighborhood C*α* RMSD measured structural deviation around mutated positions using a local sequence window around each mutation, average and minimum confidence values were parsed from predicted-structure B-factor fields when the structure-prediction workflow stored pLDDT-like confidence in that column [9].

A candidate was assigned `PASS_REFOLD_QC` when it preserved global fold geometry, active-site geometry, mutation-neighborhood geometry, and prediction confidence under the configured computational thresholds.

#### Computational admissibility criterion

Using the mutation set *M*_*k*_, the active-site reference set *A*, and the nearest active-site distance *d* (*i, A*) defined above, EditorForge summarizes final candidate admissibility as the intersection of sequence-level mask compliance, active-site-shield compliance, and structure-QC acceptance. Let *D*_shield_ denote the final active-site-shielded design mask after Active Site Shield correction, and let

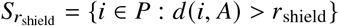

denotes the set of residue positions outside the hard active-site shield, with *r*_shield_ = 16 °A in the MMLV RT demonstration. A generated candidate sequence *s*^(*k*)^ is hard-constraint admissible only if

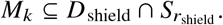

Thus, every mutation in an admissible candidate must lie both inside the corrected design envelope and outside the hard active-site shield.

For each returned predicted structure, Enhanced RefoldQC assigns a computational quality-control vector

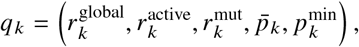

where 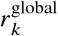 is global C*α* RMSD, 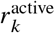 is active-site C*α* RMSD, 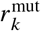 is mutation-neighborhood C*α* RMSD, 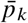 is average pLDDT-like confidence, and 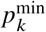 is minimum pLDDT-like confidence. Let *Q* denote the configured RefoldQC acceptance region and the final computationally accepted candidate set is therefore

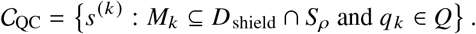

This definition is a computational triage criterion as it states that accepted candidates satisfy the corrected mutation mask, avoid hard active-site-shield breaches, and pass the configured structural-QC thresholds.

All EditorForge stages wrote machine-readable outputs, including JSON masks, CSV mutation tables, ranked candidate sheets, FASTA files, structure-prediction packets, and QC summaries. For the final preprint dataset, these outputs were compiled into a frozen evidence folder titled `Important Data`. The main candidate-level table used in the Results is TABLE 2_FINAL_CANDIDATE_DASHBOARD.csv. File-level provenance is recorded in RAW_DATA_FILE_MANIFEST.csv, and column-level metadata are recorded in `RAW_DATA_SCHEMA_DICTIONARY`.csv. Flattened JSON/JSONL outputs are provided in `ALL_JSON_JSONL_FLATTENED_ROWS_COMBINED`.csv, and parsed FASTA records are provided in `ALL_FASTA_SEQUENCE_ROWS_COMBINED`.csv. The external-source crosscheck table, `editorforge_external_online_sources_crosscheck.csv`, records the public structural, software, database, repository, and identifier sources used to ground reproducibility statements. This packaging strategy is consistent with reproducible computational reporting and FAIR data/software principles [28, 29, 30].

## 3 Results

### 3.1 EditorForge produced a conservative redesign envelope for MMLV reverse transcriptase

EditorForge was first evaluated as a constrained redesign workflow on the MMLV reverse-transcriptase parent structure. The working parent sequence contained 453 residues. Rather than permitting global redesign, EditorForge restricted ProteinMPNN generation to a 25-position mutable envelope and fixed the remaining 428 positions. Thus, only 5.5% of the parent sequence was designable, while 94.5% was required to remain parent-identical. This constraint architecture produced a deliberately conservative redesign problem as the overarching purpose was to test whether inverse-folding generation could be forced into a narrow, auditable design space rather than sequence divergence from RT parent. The fixed-position mask, active-site references, hard active-site shield, and local sequence-window protection rule together defined the allowed mutation region before candidate generation. This allowed subsequent candidate filtering to be expressed as a direct mutation-set audit: every candidate mutation either belonged to the declared design envelope or was recorded as a violation.

The initial design mask also established the active-site-protection problem that motivated the later shielded workflow. The active-site reference set contained residues 150, 224, and 225 while a hard 16 °A active-site shield and a local ± 2 sequence-neighborhood rule were applied around these reference positions. When EditorForge audited the initial design positions against this active-site protection rule, five design positions were identified as unsafe: M151, Q187, S44, Q61, and R179. Their nearest active-site-reference distances were 3.818 °A, 15.770 °A, 12.140 °A, 14.810 °A, and 11.305 °A, respectively. Logically because of this, all five were therefore inside the hard 16 °A shield and were removed from the designable set.

Active Site Shield replaced the five removed positions with lower-contact positions outside the hard shield while preserving the 25-position design budget and the replacement positions were I186, K39, R57, A296, and S116 and their nearest active-site-reference distances were 16.114 °A, 19.684 °A, 18.711 °A, 49.220 °A, and 29.422 °A, respectively. Furthermore, Their C*α* contact counts were 6, 7, 8, 11, and 11. This correction preserved the total number of designable residues while moving the mask away from active-site-proximal positions and teh final shielded design set therefore contained 25 designable positions and 428 fixed positions, but no design position inside the hard active-site shield.

### 3.2 Post-shield candidate auditing removed hard constraint violations before refolding

ProteinMPNN was rerun using the active-site-shielded mask and Post Shield Audit then evaluated 24 regenerated candidates against the corrected design envelope. In short, all 24 candidates passed the shielded constrained-design check at the mutation-integrity level: fixed-position violation count was 0 for every candidate, outside-design mutation count was 0 for every candidate, and hard active-site-shield mutation count was 0 for every candidate. The post-shield candidates carried 17-20 mutations, with a mean mutation count of 18.58 where one candidate contained 17 mutations, ten candidates contained 18 mutations, eleven candidates contained 19 mutations, and two candidaes contained 20 mutations. The nearest active-site mutation distance was 16.114 °A for all audited candidates, which placed the mutation set immediately outside the hard active-site shield, a result directly indicates that Active Site Shield removed the hard active-site-proximal failure mode observed in the initial design mask.

However, the post-shield audit retained active-site-adjacent review concerns because all 24 candidates were assigned to the `B_REVIEW_PERIPHERAL_ACTIVE_ZONE` tier because they retained mutations in the peripheral review zone outside the hard 16 °A shield, nineteen candidates had five peripheral-zone mutations, and five candidates had six peripheral-zone mutations which suggests that it cannot possibly be full biological clearance bu should be interpreted as hard-shield compliance rather than full biological clearance. EditorForge successfully removed mutations inside the prohibited active-site shield, but the peripheral-zone flags identify candidates that still require downstream structural and experimental review.

The final candidate dashboard retained the 24 audited post-shield candidates and recorded the subset with returned predicted-structure QC metrics. Eight candidates were evaluated by Enhanced RefoldQC and received PASS_REFOLD_QC: MMLV_RT_4MH8_v13_0015, MMLV_RT_4MH8_v13_0018, MMLV_RT_4MH8_v13_0005, MMLV_RT_4MH8_v13_0006, MMLV_RT_4MH8_v13_0010, MMLV_RT_4MH8_v13_0023, MMLV_RT_4MH8_v13_0008, and MMLV_RT_4MH8_v13_0002. These eight candidates had 17-18 mutations, zero fixed-position violations, zero outside-design mutations, zero hard active-site-shield breaches, and a minimum active-site mutation distance of 16.114 °A.

### 3.3 Enhanced RefoldQC showed parent-like predicted geometry for the final candidate set

Enhanced RefoldQC evaluated eight returned predicted candidate structures. Each predicted structure contained 453 C*α* atoms and was compared against the 453-residue parent structure. All eight predicted structures matched their intended design sequences with sequence identity to design equal to 1.0000. Sequence identity to the parent was 0.9625 for the 17-mutation candidate and 0.9603 for the seven 18-mutation candidates.

All eight evaluated candidates passed the configured computational refolding screen. Global C*α* RMSD values ranged from 1.2061 to 1.5555 °A, with a mean of 1.4441 °A. This indicates that the predicted structures remained close to the parent RT fold after constrained redesign. The lowest global RMSD was observed for `MMLV_RT_4MH8_v13_0008`, with a global C*α* RMSD of 1.2061 °A. The highest global RMSD was observed for `MMLV_RT_4MH8_v13_0010`, with a global C*α* RMSD of 1.5555 °A.

Active-site C*α* RMSD values ranged from 0.4098 to 1.8397 °A, with a mean of 1.5148 °A. The lowest active-site RMSD again occurred in `MMLV_RT_4MH8_v13_0008`, which had an active-site C*α* RMSD of 0.4098 °A. The largest active-site RMSD was observed for `MMLV_RT_4MH8_v13_0010`, with an active-site C*α* RMSD of 1.8397 °A. Mutation-neighborhood C*α* RMSD values ranged from 1.3155 to 1.6848 °A, with a mean of 1.5312 °A. The lowest mutation-neighborhood RMSD was again observed for `MMLV_RT_4MH8_v13_0008`, whereas the highest was observed for `MMLV_RT_4MH8_v13_0010`.

Predicted-structure confidence was consistently high across the eight evaluated candidates. Average pLDDT-like confidence values ranged from 94.87 to 95.11, with a mean of 95.04. Minimum pLDDT-like confidence values ranged from 66.50 to 72.69. These values indicate that the predicted structures used for RefoldQC were not globally low-confidence predictions, although local lower-confidence regions remained present in the minimum-confidence measurements. A normalized candidate-level QC profile for the eight `PASS_REFOLD_QC` candidates is provided in Supplementary Figure S2. This visualization summarizes relative performance across global RMSD, active-site RMSD, mutation-neighborhood RMSD, predicted confidence, and parent-sequence identity, with RMSD metrics inverted so that higher normalized values indicate stronger structural preservation.

Taken together, the results support the intended computational claim, that EditorForge constrained inverse-folding redesign to a small editor-domain design envelope, detected active-site-proximal design risk in the initial mask, corrected that risk by replacing five unsafe positions while preserving the 25-position design budget, regenerated candidates under the shielded mask, and identified eight candidates that passed computational structure-QC screening. Shortly put they show that EditorForge can convert general inverse-folding output into an auditable, active-site-aware, structure-QC-filtered candidate set.

### 3.4 Supplementary Results

## 4 Analysis and Discussion

AS stated earlier, EditorForge addresses a practical gap between inverse-folding generation and candidate selection for enzyme-like and editor-associated protein domains. Fixed-backbone inverse-folding methods can rapidly propose sequences compatible with a supplied structure, but structural compatibility is not equivalent to design suitability [6, 4]. For catalytic or editor-associated proteins, a generated sequence can preserve a plausible global fold while still placing mutations near catalytic, substrate-proximal, conserved, or structurally sensitive regions. In these cases, the central design question extends beyond backbone compatibility to a declared set of biological and structural constraints.

The MMLV reverse-transcriptase demonstration shows how this kind of controlled triage can be implemented in a reproducible way. As shown, EditorForge restricted redesign to 25 positions in a 453-residue parent domain and fixed the remaining 428 positions in order to test whether inverse-folding generation could be restricted to a narrow and auditable region of sequence space. In this framing, ProteinMPNN can be thought of as a generator while EditorForge enforces the admissible candidate class. A useful way to describe the workflow is as a sequence of filters applied to raw inverse-folding output. If we let *s*^(0)^ denote the parent sequence, let *P* = {1, …, *N*} be the residue index set, and let *D* ⊂ *P* be the designable set, for a generated candidate *s*^(*k*)^, the mutation set is

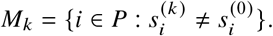

The first constraint is simply means that mutations must remain inside the declared design envelope:

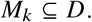

This condition may appear basic, but is very important since it distinguishes raw inverse-folding output from candidates that actually obey the intended redesign problem.

However, the study shows that this condition alone is not sufficient for editor-domain redesign due to the fact that a mutation can be formally allowed by the design mask while still being too close to a catalytic-reference region. This is the failure mode that motivated Active Site Shield. For the MMLV RT demonstration, the active-site reference set was defined as

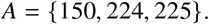

Given a hard shield radius *r*_shield_ = 16 °A, the safe region can be understood as the set of residue positions whose nearest active-site-reference distance exceeds the cutoff. Thus, the hard-shield rule adds a second admissibility condition:

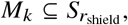

where 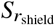 denotes positions outside the hard active-site shield. A candidate that satisfies both the design-mask rule and the hard-shield rule therefore lies in the intersection

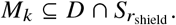

This compact expression captures the central logic of the workflow where the effective design space is the subset of those positions that also respects active-site geometry.

The initial design mask violated this stricter interpretations as five positions in the initial 25-position envelope fell inside the hard active-site shield: M151, Q187, S44, Q61, and R179. Their nearest active-site-reference distances were 3.818 °A, 15.770 °A, 12.140 °A, 14.810 °A, and 11.305 °A, respectively. These positions were not ordinary outside-design mutations, because they belonged to the declared designable set so the error was instead in the design space itself: the initial mask included positions that were designable by the mask but unsafe by the active-site rule, revealing the crutical fact that fixed-position enforcement can appear successful while the design problem remains biologically risky.

Active Site Shield corrected this by removing unsafe design positions and replacing them with lower-contact alternatives outside the hard shield. The five replacement positions were I186, K39, R57, A296, and S116, with nearest active-site-reference distances of 16.114 °A, 19.684 °A, 18.711 °A, 49.220 °A, and 29.422 °A, respectively. The key impact is that this particular module changed the allowed design space and then required regeneration under the corrected mask. In terms of workflow logic, EditorForge implements the continous loop of mask construction, candidate generation, audits, mask correction, regeneration, then repeat. This is one of the main methodological contributions of the work. The before/after correction is shown in Figure 3.

**Figure 1:**
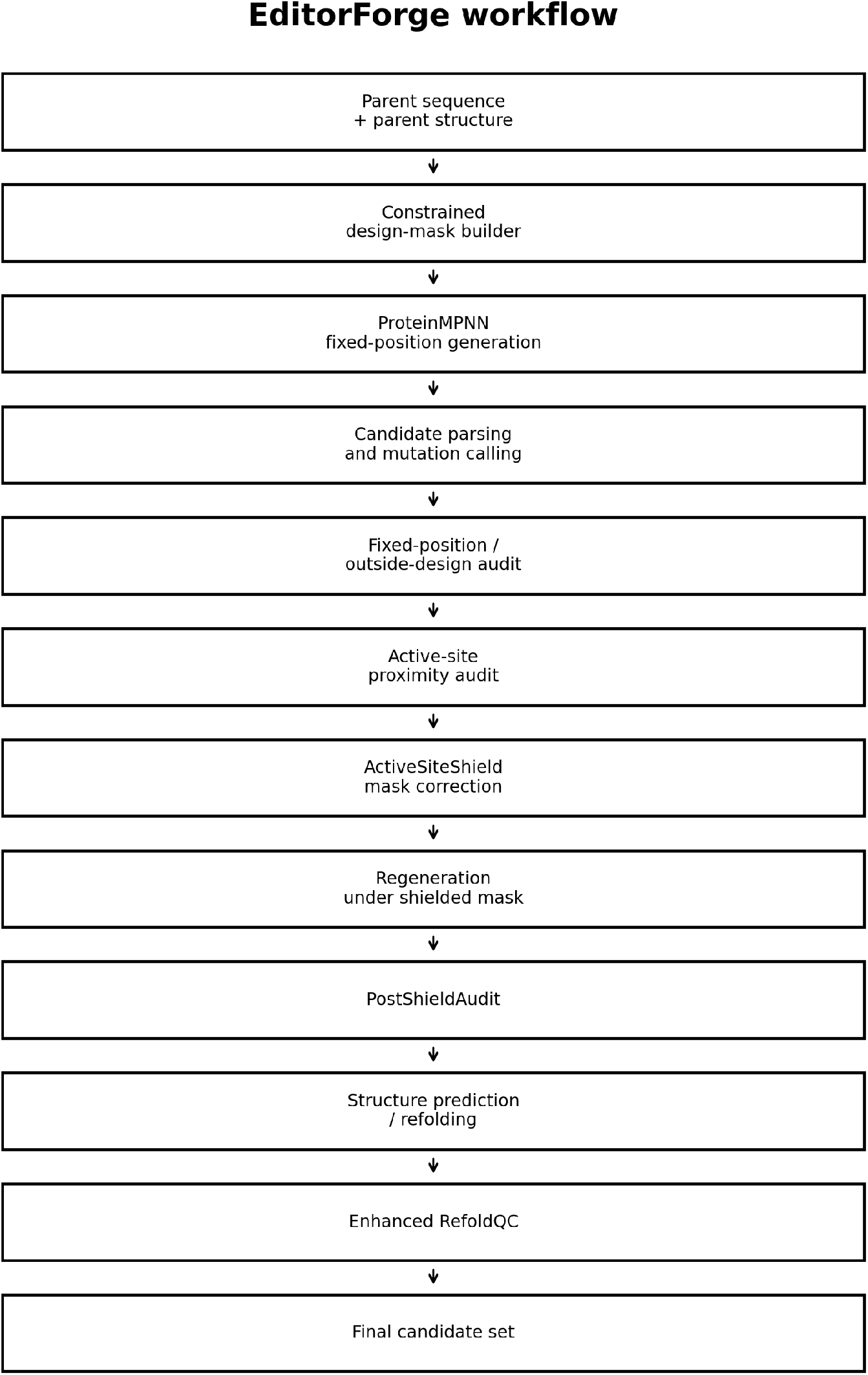
EditorForge workflow for constrained inverse-folding triage of editor-associated protein domains. EditorForge begins with a parent sequence and parent structure, constructs a residue-level design mask, runs ProteinMPNN under fixed-position constraints, parses generated candidates, audits mutations against the allowed design envelope, evaluates active-site-proximal risk, applies Active Site Shield when unsafe design positions are detected, regenerates candidates under the corrected shielded mask, and evaluates final predicted structures with Enhanced RefoldQC. The workflow is designed to sit between inverse-folding generation and downstream validation since it provides a reproducible constraint-and-audit layer that converts general inverse-folding output into a smaller set of candidate designs with explicit mask compliance, active-site-shield compliance, and structure-QC records.

**Figure 2:**
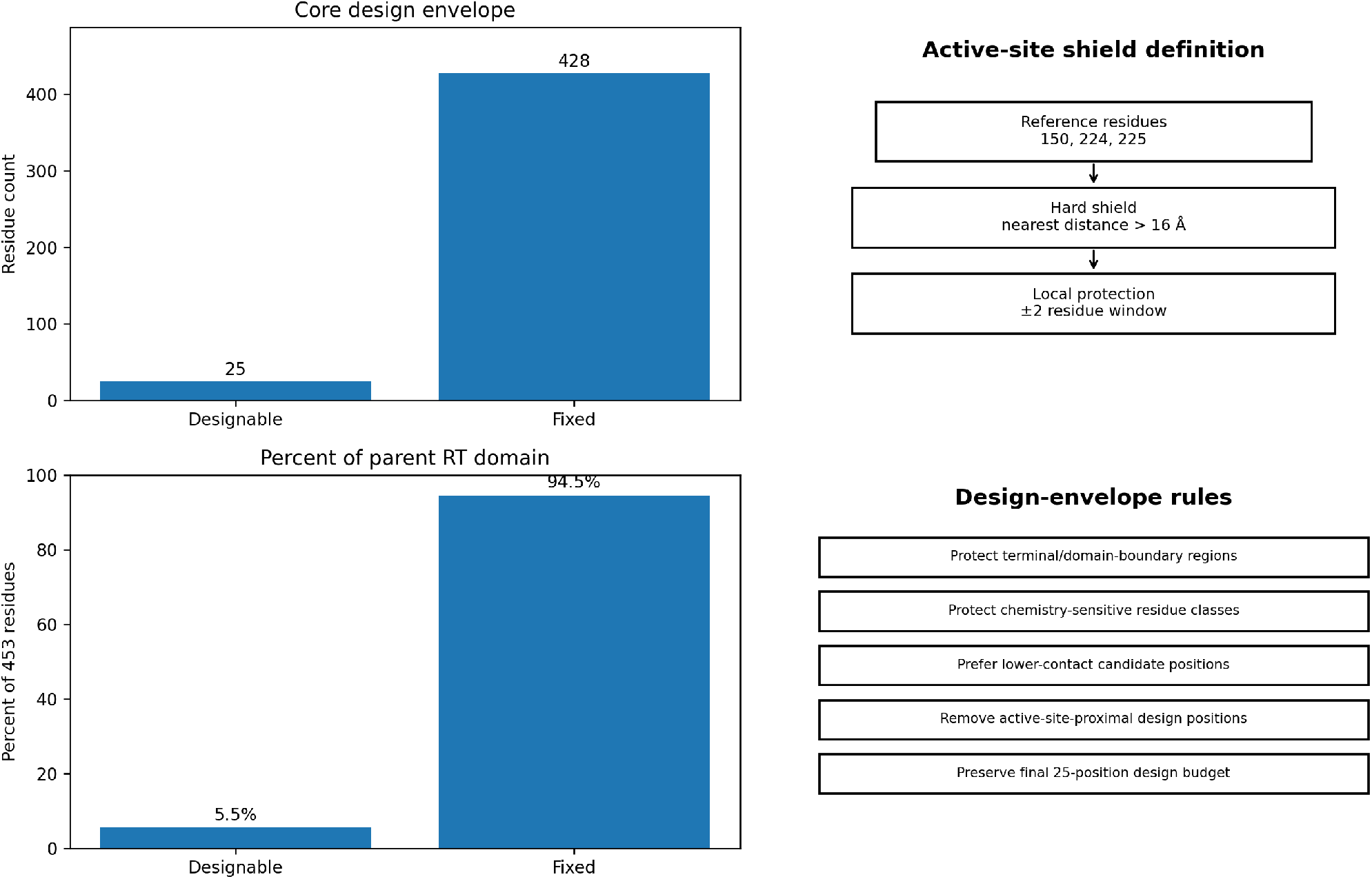
Constraint architecture for the MMLV reverse-transcriptase demonstration target. The working MMLV RT parent contained 453 residues. EditorForge restricted inverse-folding redesign to a conservative 25-residue design envelope while fixing the remaining 428 residues as parent-identical positions. Thus, only 5.5% of the parent RT domain was designable, whereas 94.5% was fixed. The active-site protection layer used reference residues 150, 224, and 225, a hard nearest-distance shield of 16 °A, and a local ± 2 residue sequence-window rule. Additional constraint rules protected terminal and domain-boundary regions, chemistry-sensitive residues, and high-contact structural positions while favoring lower-contact alternatives when replacement design positions were needed. This architecture defines the residue-level rules that constrained ProteinMPNN generation before candidate auditing and active-site shielding.

**Figure 3:**
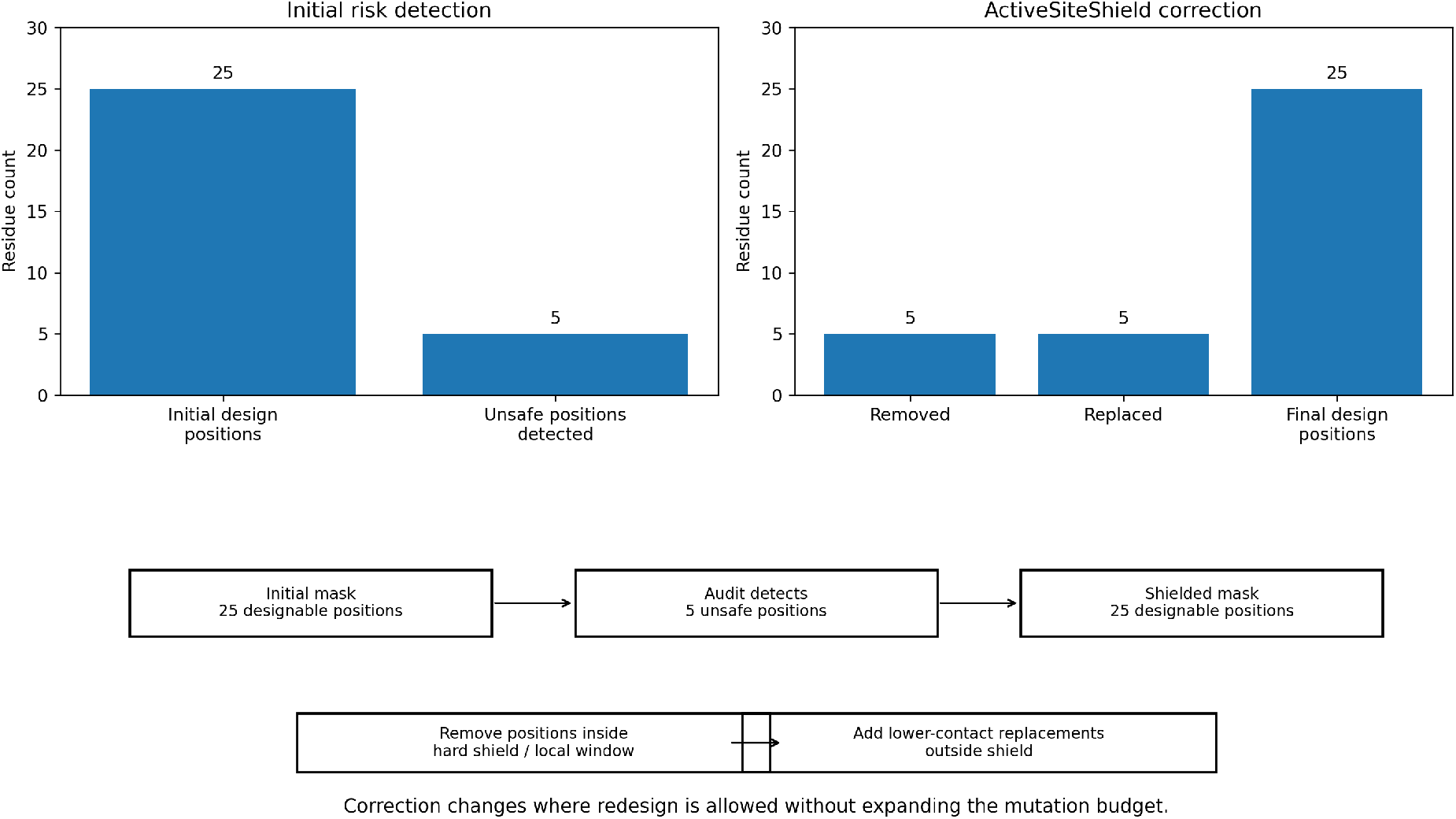
Active Site Shield correction of active-site-proximal design positions. The initial 25-position design envelope contained five positions inside the hard active-site shield: M151, Q187, S44, Q61, and R179. Their nearest active-site-reference distances were 3.818 °A, 15.770 °A, 12.140 °A, 14.810 °A, and 11.305 °A, respectively, placing all five within the 16 °A exclusion rule around active-site reference residues 150, 224, and 225. Active Site Shield removed these unsafe positions and added five lower-contact replacement positions outside the hard shield: I186, K39, R57, A296, and S116. The correction preserved the 25-position design budget while moving the allowed mutation envelope away from active-site-proximal residues. This result demonstrates that merely than re-auditing inverse folding outputs, EditorForge can revise the design mask and regenerate candidates under a corrected constraint architecture.

The post-shield audit showed that this correction removed the hard active-site-proximal failure mode. After regeneration under the shielded mask, 24 post-shield candidates were audited where all 24 had zero fixed-position violations, zero outside-design mutations, and zero hard active-site-shield breaches. These results show that the final shielded workflow produced candidates satisfying the intended sequence-level and hard-geometric rules. Of course, the candidate funnel and audit matrix are shown in Figure 4, and the final candidate-level dashboard is reported in Table 2.

**Figure 4:**
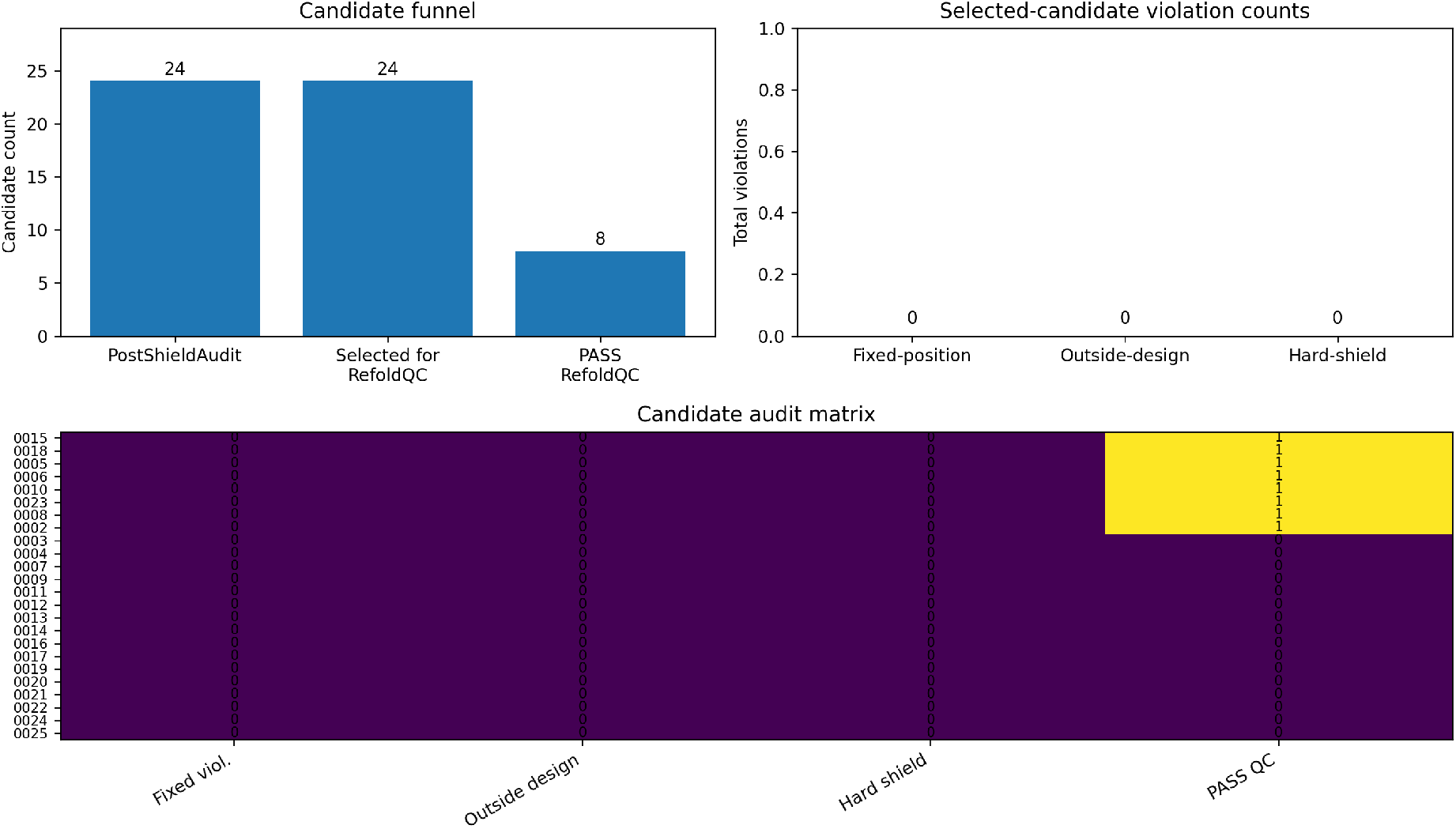
Post-shield candidate funnel and mutation-integrity audit. After Active Site Shield correction, ProteinMPNN was rerun under the shielded 25-position design mask and 24 regenerated candidates were audited. Post Shield Audit found zero fixed-position violations, zero outside-design mutations, and zero hard active-site-shield breaches across the audited post-shield set. The final candidate dashboard retained the 24 post-shield candidates and identified eight candidates with returned predicted structures that passed Enhanced RefoldQC. All eight passing candidates had 17-18 mutations, no hard-shield breaches, and a nearest active-site mutation distance of 16.114 °A. The audit matrix separates hard constraint compliance from downstream review: the candidates satisfy the hard shield, but peripheral-zone flags remain a reason for structural and biological follow-up.

At the same time, the post-shield candidates retained peripheral-zone flags. For exampe, a hard shield creates a prohibited region, but it does not mean every position outside that prohibited region is automatically biologically neutral. Mutations outside the hard shield may still be close enough to functional regions to require review. In the post-shield audit, all 24 candidates were assigned to the `B_REVIEW_PERIPHERAL_ACTIVE_ZONE` tier, while still having zero mutations inside the hard shield. EditorForge therefore distinguishes between hard violations and softer proximity concerns, appropriate for editor-associated domains, where a computational workflow should remove clear failures without pretending that all remaining candidates are experimentally cleared.

Enhanced RefoldQC adds a third layer of candidate selection. After sequence-mask compliance and active-site-shield compliance, the returned predicted structures were evaluated for parent-like geometry. Each candidate can be summarized by a QC vector,

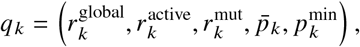

where 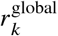 is global C*α* RMSD, 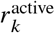 s active-site C*α* RMSD, 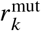 is mutation-neighborhood C*α* RMSD, 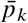 is average pLDDT-like confidence, and 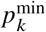 is minimum pLDDT-like confidence. This vector is useful because it separates different dimensions of structural plausibility because of the fact that a candidate can preserve the global fold while showing more local deviation near the active site or mutation neighborhood while conversely, a candidate can have acceptable local geometry while showing broader structural drift.

In this study, eight returned predicted structures passed the configured RefoldQC screen. Their global C*α* RMSD values ranged from 1.2061 to 1.5555 °A, active-site C*α* RMSD values ranged from 0.4098 to 1.8397 °A, and mutation-neighborhood C*α* RMSD values ranged from 1.3155 to 1.6848 °A. Also average pLDDT-like confidence values ranged from 94.87 to 95.11, while minimum pLDDT-like confidence values ranged from 66.50 to 72.69. The final computationally accepted set can therefore be described as candidates satisfying three conditions: mask compliance, hard-shield compliance, and RefoldQC acceptance or the final evaluated set is

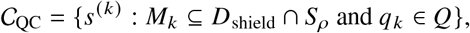

where *D*_shield_ is the corrected design mask and *Q* is the configured RefoldQC acceptance region. This expression is useful because it identified candidates that passed a declared computational admissibility test.

The strongest candidate-level structural result was observed for `MMLV_RT_4MH8_v13_0008`. This candidate had the lowest global C*α* RMSD, the lowest active-site C*α* RMSD, and the lowest mutation-neighborhood C*α* RMSD among the eight passing candidates, with values of 1.2061 °A, 0.4098 °A, and 1.3155 °A, respectively. Conversely, `MMLV_RT_4MH8_v13_0010` had the highest values across those three RMSD categories among the passing set, with values of 1.5555 °A, 1.8397 °A, and 1.6848 °A. This does not prove that v13 0008 is functionally superior or that `v13_0010` is functionally inferior; instead this only provides a structural-prioritization basis for downstream review onwards. In an experimental pipeline, `v13_0008` would be a natural first candidate to inspect structurally in more detail because it was the most parent-like by the measured RefoldQC geometry. The complete RefoldQC metric distribution is shown in Figure 5.

**Figure 5:**
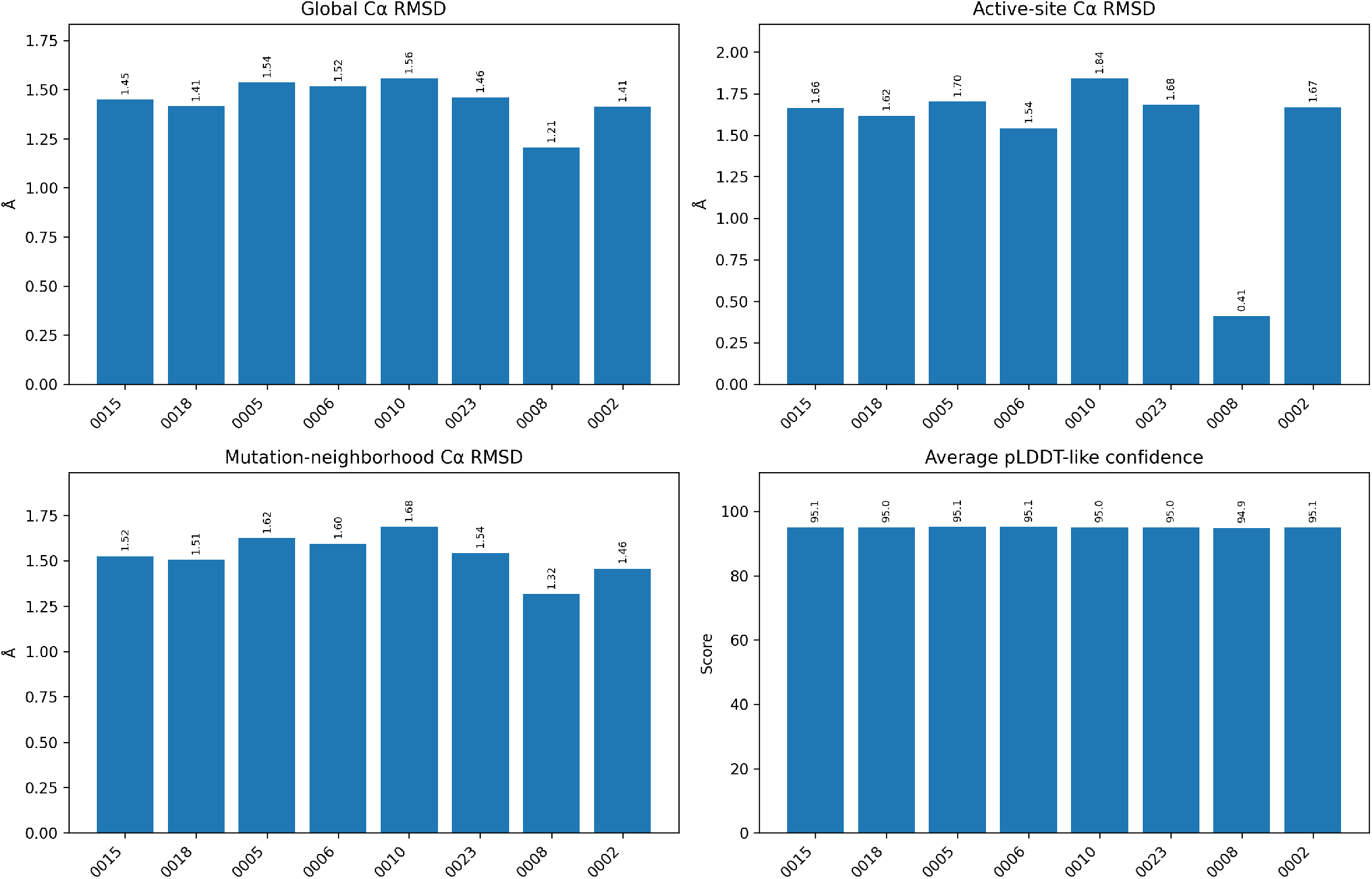
Enhanced RefoldQC metrics for the eight final passing candidates. Enhanced RefoldQC compared each returned predicted candidate structure against the parent MMLV RT structure using global C*α* RMSD, active-site C*α* RMSD, mutation-neighborhood C*α* RMSD, and average pLDDT-like confidence. All eight evaluated candidates received `PASS_REFOLD_QC`. Global C*α* RMSD values ranged from 1.2061 to 1.5555 °A, active-site C*α* RMSD values ranged from 0.4098 to 1.8397 °A, and mutation-neighborhood C*α* RMSD values ranged from 1.3155 to 1.6848 °A. Average pLDDT-like confidence values ranged from 94.87 to 95.11. These results support the computational conclusion that the final candidates preserved parent-like predicted geometry under the configured RefoldQC screen. They do not establish improved enzymatic activity, editing performance, fidelity, expression, delivery, or safety.

**Figure 6:**
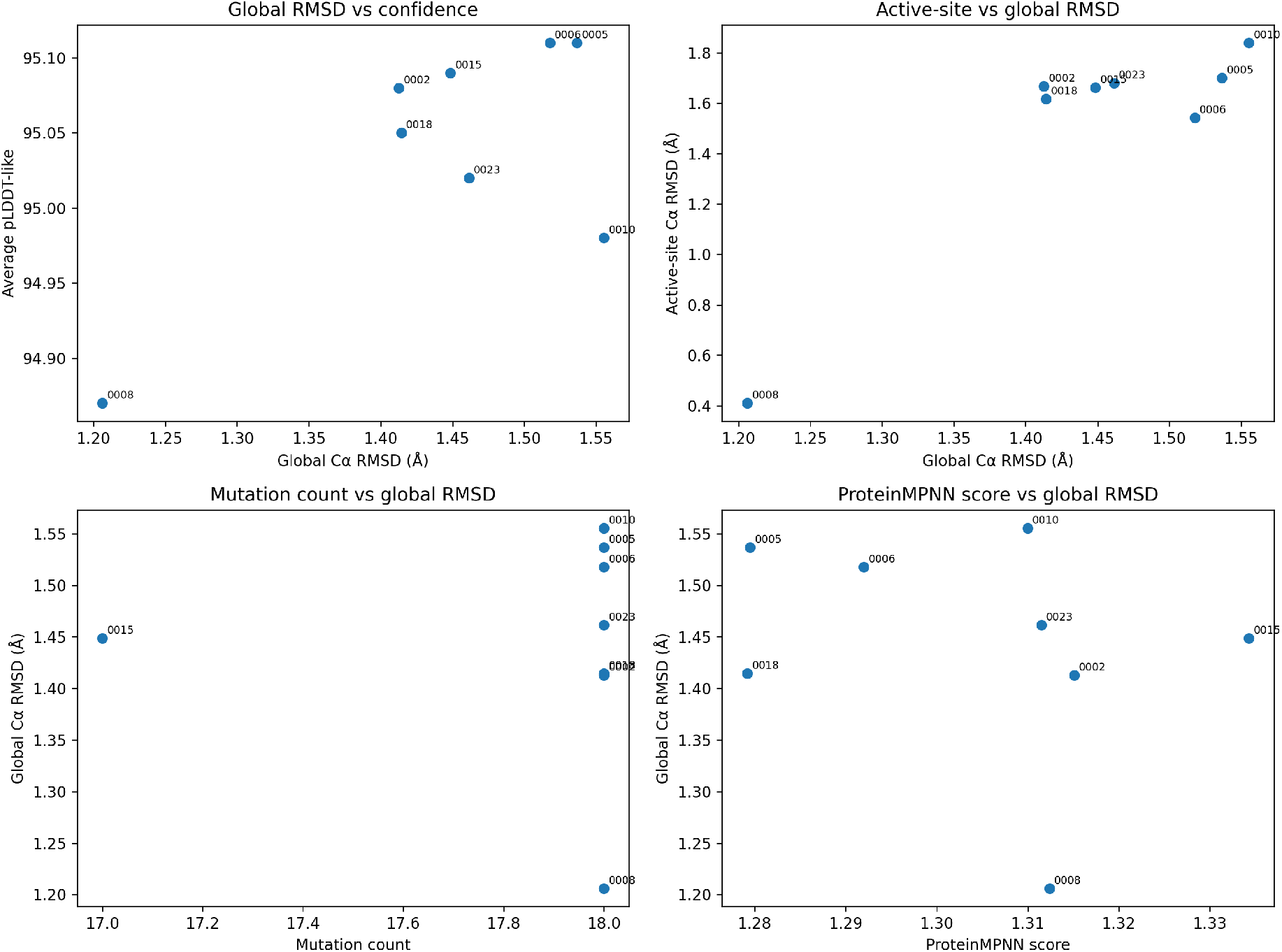
Relationships among generation and RefoldQC metrics for evaluated EditorForge candidates. Scatter plots compare global C*α* RMSD against average pLDDT-like confidence, active-site C*α* RMSD against global C*α* RMSD, mutation count against global C*α* RMSD, and ProteinMPNN score against global C*α* RMSD. These diagnostic relationships are intended to expose obvious tradeoffs between mutation burden, inverse-folding score, predicted-structure confidence, and geometric deviation from the parent structure. In the final evaluated set, the passing candidates occupied a narrow structural-QC range, with global C*α* RMSD values between 1.2061 and 1.5555 °A and average pLDDT-like confidence values between 94.87 and 95.11. However, this figure is a computational diagnostic rather than evidence of functional improvement because at n=8 correlations are often weak

**Figure 7:**
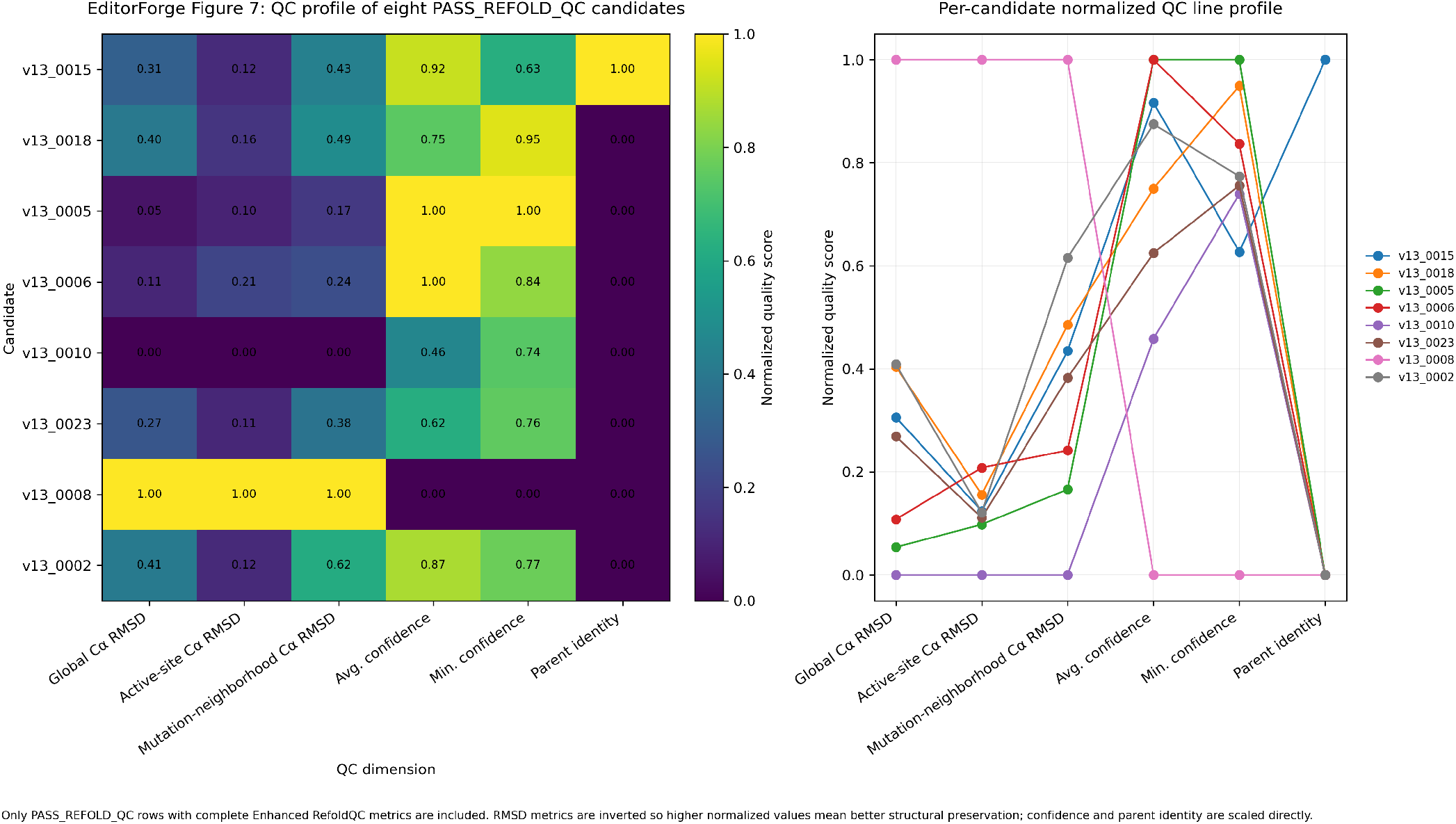
Normalized candidate-level QC profile for the eight PASS REFOLD QC candidates. The heatmap and line profile summarize normalized quality-control behavior across the eight candidates that passed Enhanced RefoldQC. The plotted dimensions are global C*α* RMSD, active-site C*α* RMSD, mutation-neighborhood C*α* RMSD, average pLDDT-like confidence, minimum pLDDT-like confidence, and parent-sequence identity. RMSD metrics are inverted before normalization so that higher normalized values indicate stronger structural preservation, while confidence and parent-identity values are scaled directly. Values are normalized within the eight-candidate passing set and therefore represent relative QC profiles, not independent biological scores. This visualization supports candidate-level comparison across multiple structural-QC dimensions.

**Figure 8:**
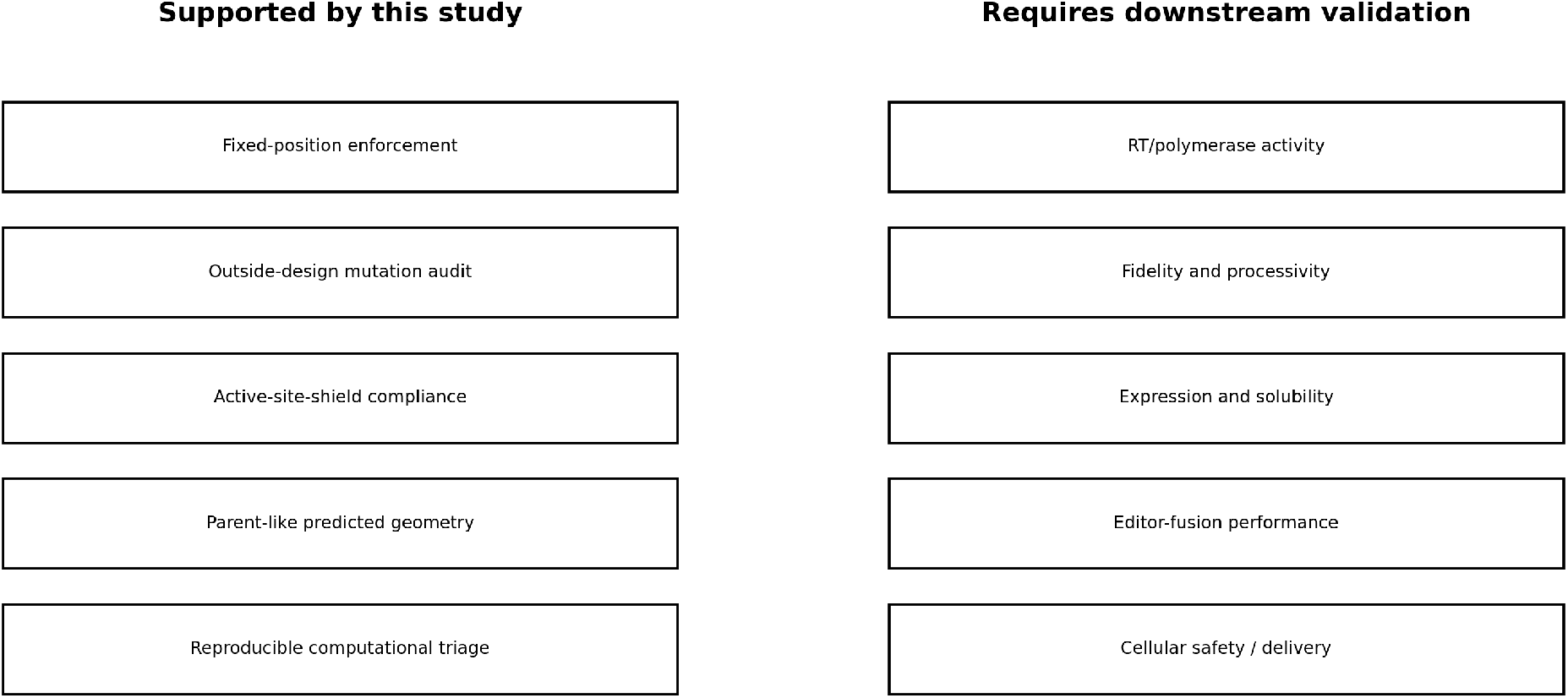
Boundary between EditorForge-supported computational claims and downstream biological validation requirements. EditorForge supports constrained computational triage claims: fixed-position enforcement, outside-design mutation auditing, active-site-shield compliance, parent-like predicted geometry, and reproducible candidate filtering. These outputs justify advancing selected candidates to further review, but they do not establish improved reverse-transcriptase activity, fidelity, processivity, expression, solubility, editor-fusion performance, cellular delivery, or biological safety. The right-hand side lists validation categories that remain outside the scope of this study and require downstream biochemical, biophysical, structural, or cellular assays. This boundary is central to the interpretation of the work: `PASS_REFOLD_QC` is a computational triage label, not an experimental function label.

The study also illustrates why multiple computational metrics should not be collapsed into a single design score because as seen ProteinMPNN score, mutation count, fixed-position integrity, outside-design status, active-site distance, peripheral-zone status, global RMSD, local RMSD, and predicted confidence each measure different aspects of candidate quality. For example, ProteinMPNN score reflects compatibility with the supplied backbone under the inverse-folding model. Mutation audits measure obedience to the declared sequence-design problem while active-site distances measure compliance with a functional-geometry rule and RefoldQC RMSD values measure predicted structural preservation. Furthermore confidence scores indicate whether the predicted structures are reliable enough to support geometric comparison, so treating these separately ensures scientific integrity. Also, this separation is especially important for editor-associated proteins because genome editors, reverse transcriptases, nucleases, deaminases, recombinases, polymerases, and related enzyme-like domains often depend on localized structural features [14, 15, 23, 22]. Catalytic residues, metal-coordination environments, nucleic-acid-contact surfaces, substrate-positioning loops, domain interfaces, and conformational transitions can be disrupted even when a predicted global fold appears preserved. Thus, EditorForge is built around this problem in order to reduce the chance that constraint-violating or active-site-proximal candidates are advanced as serious designs.

The workflow also shifts emphasis from candidate ranking to design-space control as many computational design pipelines generate many candidates and then select the best-scoring outputs. Instead, EditorForge asks whether the candidate-generation problem was properly constrained before ranking occurs. Ultimately, this is important because a high-scoring sequence generated from an unsafe design space can still be inappropriate for an editor-domain application. In the MMLV RT demonstration, the initial mask required correction before the final candidate set could be interpreted as active-site-aware.

Tracebility is also another important fature, as each stage of EditorForge produces machine-readable output: design masks, mutation tables, audit reports, candidate FASTA files, structure-prediction packets, RefoldQC summaries, file manifests, schema dictionaries, and final candidate dashboards. This makes the path from parent structure to final candidate table reconstructable. The frozen evidence package includes RAW_DATA_FILE_MANIFEST.csv, `RAW_DATA_SCHEMA_DICTIONARY.csv, ALL_JSON_JSONL_FLATTENED_ROWS_COMBINED.csv, and ALL_FASTA_SEQUENCE_ROWS_COMBIN` These files document raw source files, schema metadata, flattened JSON/JSONL outputs, and parsed sequence records, respectively. This transparency also prevents a common weakness in presenting only attractive final candidates while leaving the filtering path unclear. To combat this, EditorForge makes reports that the initial mask contained unsafe positions, that Active Site Shield corrected the mask, that post-shield candidates retained peripheral review flags, and that RefoldQC was performed on the returned predicted structures.

Overall, EditorForge converted fixed-backbone inverse-folding output into a constrained, active-site-aware, structurally triaged candidate set for an editor-associated RT domain as it restricted generation to a 25-position design envelope, detected five active-site-proximal design positions, replaced those five positions while preserving the 25-position design budget, regenerated 24 candidates under stricter constraints, confirmed zero hard mutation-integrity violations in the audited post-shield set, and identified eight returned predicted structures that passed RefoldQC. This conclusion is summarized in Table 1 and Table 3.

**Table 1:** EditorForge pipeline milestones and quantitative outputs for the MMLV RT demonstration. This table summarizes the major computational stages of the EditorForge workflow and the specific question answered by each stage. The pipeline first restricted ProteinMPNN generation to a 25-position design envelope while fixing 428 residues, then used ActiveSiteShield to remove five active-site-proximal design positions and add five safer replacement positions. PostShieldAudit retained 24 regenerated candidates with zero fixed-position violations, zero outside-design mutations, and zero hard active-site-shield breaches. Enhanced RefoldQC was then applied to eight returned predicted candidate structures, all of which received `PASS_REFOLD_QC` status. These milestones define the computational evidence chain from constrained generation to final structural triage.

| Stage | Question answered | Quantitative result | Manuscript role |
| --- | --- | --- | --- |
| Constrained design setup | Can inverse folding be restricted to a small editor-domain design envelope? | 25 designable residues; 428 fixed residues; 453 total residues | Defines the conservative redesign problem |
| Initial active-site audit | Did the initial design mask contain active-site-proximal risk? | 5 unsafe design positions detected inside the 16 Å hard shield | Motivates ActiveSiteShield |
| ActiveSiteShield correction | Can unsafe design positions be removed while preserving the design budget? | 5 unsafe positions removed; 5 replacements added; 25-position budget preserved | Corrects the design mask |
| PostShieldAudit | Did regenerated candidates obey the corrected hard constraints? | 24 candidates audited; 0 fixed-position violations; 0 outside-design mutations; 0 hard-shield breaches | Confirms hard constraint compliance before structure QC |
| Enhanced RefoldQC | Did returned predicted structures preserve parent-like geometry? | 8 returned predicted structures evaluated; 8 PASS_REFOLD_QC | Final computational structure-QC result |

**Table 2:**
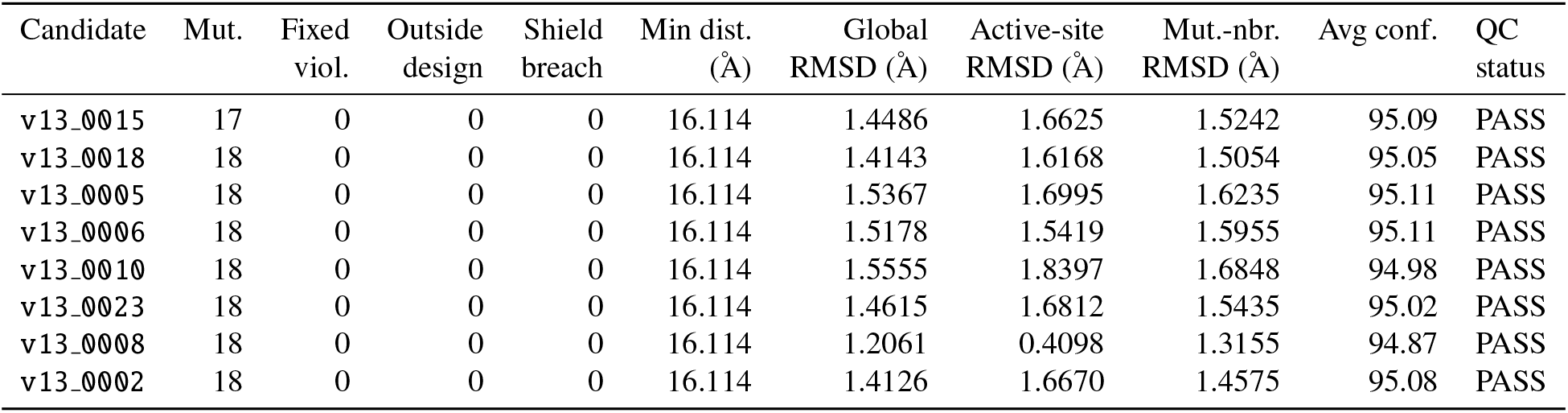
Final candidate dashboard for the eight EditorForge candidates that passed Enhanced RefoldQC. The table reports mutation count, fixed-position violations, outside-design mutations, hard active-site-shield breaches, nearest active-site mutation distance, global C*α* RMSD, active-site C*α* RMSD, mutation-neighborhood C*α* RMSD, average pLDDT-like confidence, minimum pLDDT-like confidence, and QC status. All eight candidates had zero fixed-position violations, zero outside-design mutations, zero hard active-site-shield breaches, and `PASS_REFOLD_QC` status.

**Table 3:** Supported claims and nonclaims for the EditorForge MMLV RT demonstration. The supported claims are restricted to computational triage: constrained inverse-folding design, active-site-aware mask correction, and structural preservation under RefoldQC. The nonclaims identify biological conclusions that are not supported by the current dataset because no wet-lab validation was performed. This table complements Figure 8.

| Claim | Supported by | Status |
| --- | --- | --- |
| EditorForge constrains inverse-folding design to an editor-domain design envelope | 25 designable positions; 428 fixed positions | Supported computationally |
| ActiveSiteShield detects and corrects active-site-proximal design failures | 5 unsafe design positions removed and 5 replacements added | Supported computationally |
| Final candidates preserve global and local geometry under refold QC | 8/8 PASS_REFOLD_QC | Supported computationally |
| Candidates improve RT/editor function | No wet-lab assay in current work | Not claimed |
| Candidates improve fidelity, processivity, expression, or safety | No experimental validation in current work | Not claimed |

This boundary is important because the practical value of EditorForge lies in disciplined triage. This is because Wet-lab validation is expensive and capacity-limited so a computational workflow that eliminates obvious constraint failures and produces a small, auditable, structurally plausible candidate set can be useful even before experimental validation. Additionally, future versions of EditorForge could extend this framework in several directions.

First, active-site shielding could be made state-aware by using multiple parent structures, ligand-bound structures, nucleic-acid-bound structures, metal-bound structures, or predicted conformational ensembles [13]. Second, replacement-position selection could include additional priors such as solvent exposure, evolutionary conservation, coevolutionary coupling, side-chain environment, or predicted local frustration. Third, RefoldQC could be expanded beyond C*α* RMSD to include all-atom relaxation, side-chain rotamer quality, hydrogen-bond preservation, electrostatic compatibility, nucleic-acid-interface geometry, molecular-dynamics stability, or ensemble-based scoring. Fourth, the workflow could be tested across multiple editor-associated scaffolds to evaluate whether the mask-audit-shield-regenerate pattern generalizes beyond MMLV RT. In addition, EditorForge accepts sequence proposals from ProteinMPNN, BC-DESIGN, ESM-IF, or other inverse-folding engines, then applies target-specific constraint auditing and structural QC [4, 5, 3].

The most direct next step is to test whether EditorForge improves downstream candidate quality relative to less constrained inverse-folding workflows. For example a future comparison could generate one candidate set using ordinary fixed-position inverse folding and another using EditorForge with active-site shielding and RefoldQC. The two sets could then be compared experimentally for expression, solubility, stability, activity retention, fidelity, or editor compatibility. Such a study would test whether EditorForge is reproducible and whether its constraints improve the hit rate of candidates entering experimental validation.

In summary, EditorForge provides a reproducible constraint-and-audit framework for editor-domain inverse folding. The MMLV RT demonstration shows that ordinary fixed-position enforcement can miss active-site-proximal design-space risk, that this risk can be corrected by revising the design mask, and that regenerated candidates can be structurally triaged by RefoldQC. At the end of the day, this computational triage produced an auditable candidate set that obeyed the corrected design envelope, avoided hard active-site-shield breaches, and preserved parent-like predicted geometry among the evaluated predicted structures, but whether this would hold in wet-lab settings is a different situation that requires future work and proofing to be sufficient.

## 5 Limitations and nonclaims

This study has several limitations that define the scope of the present EditorForge demonstration:

The workflow was evaluated on a single editor-associated protein domain, MMLV reverse transcriptase, using PDB structure 4MH8 as the parent structural model [16, 17]. Although this is enough to show that the pipeline can be executed end to end, from target preparation through mask construction, inverse-folding generation, active-site shielding, candidate auditing, and RefoldQC triage, it does not, by itself however, establish that the same parameter choices or design rules will transfer unchanged to nucleases, deaminases, recombinases, polymerase variants, or multi-domain editor fusions. Those systems may have different domain boundaries, functional surfaces, catalytic geometries, conformational states, substrate-contact regions, metal-binding environments, or allosteric coupling patterns, and each of those differences could change how the design envelope should be built.

The active-site protection layer should also be interpreted as a demonstration-specific structural rule. In this study, the active-site-reference set was

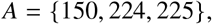

and the hard shield cutoff was *r*_shield_ = 16 °A. These values define the exclusion geometry used for the MMLV RT run rather than universal biological constants. A different parent structure, ligand-bound state, substrate-bound state, metal-bound state, or engineering objective could require different reference residues, different shield distances, or multiple protected functional surfaces. The same caution applies to the broader mask-building logic. Terminal protection, chemistry-sensitive residue protection, C*α* contact-count filtering, and lower-contact replacement selection reduce obvious redesign risk, but they remain structural heuristics. Some low-contact residues may still be functionally important, some high-contact residues may tolerate substitution, and combinations of individually reasonable substitutions may interact through epistasis. A related limitation is that the parent structure is treated as the main geometric reference. This is appropriate for a fixed-backbone inverse-folding workflow, especially in an initial computational triage setting, although reverse transcriptases and other editor-associated proteins are not static objects in their biochemical context [22, 23, 15]. For instance, substrate binding, catalysis, primer-template positioning, metal coordination, domain motion, and editor-fusion architecture can all change which residues matter. A position that appears distant from the chosen reference residues in one structure may become more important in another state, while a position that appears geometrically close in one static model may not carry the same functional risk under every biochemical condition. For this reason, hard-shield compliance is defined here as a geometric screen: it confirms that mutations avoided the prohibited active-site shell used in this study, while leaving finer functional interpretation to later structural and experimental review.

Enhanced RefoldQC provides another useful but bounded layer of evidence. The reported global C*α* RMSD, active-site C*α* RMSD, mutation-neighborhood C*α* RMSD, and pLDDT-like confidence values describe predicted structural preservation under the configured computational screen [24, 25, 26, 27]These quantities are useful for detecting large predicted disruptions in the parent fold or in local regions of concern. They do not measure expression, solubility, aggregation, proteolytic stability, enzymatic activity, fidelity, processivity, delivery, immunogenicity, cellular safety, or editor-fusion performance. C*α* RMSD is also a coarse backbone-level metric, so all-atom packing, side-chain rotamers, hydrogen-bond networks, electrostatics, metal coordination, substrate positioning, and catalytic transition-state geometry remain outside the current RefoldQC readout.

High average pLDDT-like confidence supports the use of the predicted structures for geometric comparison, but it should not be mistaken for evidence that the same conformations are experimentally populated or functionally productive [9, 10, 11]. The present evidence package is computational, retrospective, and limited in candidate number. Eight returned predicted structures passed Enhanced RefoldQC, which is sufficient for demonstrating the workflow but too small for learning general relationships among ProteinMPNN score, mutation count, RMSD, confidence, and biological viability. The scatter plots and dashboards are diagnostics for this run, not general threshold rules for editor-domain redesign. EditorForge also inherits assumptions from the external inverse-folding and structure-prediction tools that provide its inputs. ProteinMPNN supplies the sequence-generation step, while predicted structures may come from ColabFold, AlphaFold-style workflows, ESMFold, or related predictors [4, 9, 10, 11]. If those models introduce training-distribution bias, overconfident structural predictions, or systematic errors for a particular scaffold, EditorForge can audit and constrain their outputs but cannot remove every model-level failure mode. In a way, the central nonclaim of this preprint is simply just just because a candidate obtained PASS REFOLD QC, it does not indicate functional improvement.

Despite these limitations, the workflow provides a useful foundation for future work. The immediate next step is to test whether the same constraint-and-audit logic generalizes across additional editor-associated domains. A second computational target would help determine whether the ActiveSiteShield correction pattern is specific to MMLV RT or reflects a broader design-space failure mode in editor-domain inverse folding. A third target could test whether the workflow can handle different functional geometries, such as nucleic-acid-binding surfaces, metal-dependent catalytic centers, or multi-domain interfaces.

The next experimental step would be to select a small number of candidates for physical testing. Based on the current RefoldQC results, candidates with lower global and local RMSD values could be prioritized for expression and stability assays. If expression is successful, reverse-transcriptase activity, fidelity, and processivity could be evaluated in vitro. If the candidates are intended for genome-editor architectures, editor-fusion performance and cellular assays would then be required. These experiments would not simply validate the candidates; they would also validate whether EditorForge triage meaningfully improves the quality of candidates entering the experimental pipeline. In summary, the MMLV RT demonstration supports that EditorForge can detect unsafe design-space choices, correct the design mask, regenerate candidates under stricter constraints, and identify structurally plausible candidates for downstream validation.

## 6 Data availability and external-source crosscheck

All EditorForge outputs used for this preprint were compiled into a frozen evidence package before final manuscript assembly that can be found on GitHub repository: https://github.com/anthonycehn-eng/EditorForge-An-Active-Site-Aware-Framework-for-Inverse-Folding-Based-Protein-Redesign.git. The purpose of this package is to make the computational workflow traceable from the public parent structure through design-mask construction, ProteinMPNN generation, Active Site Shield correction, post-shield candidate auditing, structure-prediction handoff, Enhanced RefoldQC, manuscript tables, and final figure generation [28, 29, 30].The repository therefore serves as both a code archive and the evidence record supporting the candidate-level claims in this study.

The most important manuscript-facing table is TABLE 2 FINAL_CANDIDATE_DASHBOARD.csv because this table supports the main Results claims by recording candidate IDs, mutation counts, fixed-position violations, outside-design mutations, hard active-site-shield breaches, nearest active-site mutation distances, RMSD metrics, pLDDT-like confidence values, and RefoldQC status. It is the only raw-style candidate table that should appear directly in the main paper, because it links the final eight passing candidates to the numerical claims reported in the Results section.

The complete file-level provenance record is provided as RAW_DATA_FILE_MANIFEST.csv. This manifest records the raw files included in the evidence package, their source paths, copied filenames, file sizes, checksums, parse status, and provenance information. It is intended to make the evidence folder auditable rather than merely descriptive. Also the accompanying `RAW_DATA_SCHEMA_DICTIONARY`.csv provides column-level metadata for the raw and compiled tables, including inferred data types and example values. Together, these two files document what data exist, where the data came from, and how the tabular outputs should be interpreted.

Two combined supplementary data tables capture broader machine-readable output from the workflow.

`ALL_JSON_JSONL_FLATTENED_ROWS_COMBINED`.csv contains flattened JSON and JSONL rows from masks, audit objects, manifests, candidate-intelligence outputs, and QC summaries.`ALL_FASTA_SEQUENCE_ROWS_COMBINED`.csv contains parsed parent and candidate FASTA records, sequence lengths, sequences, and amino-acid count columns. These files are not intended to be printed in the main manuscript. They are repository-level data products that allow downstream users to inspect the sequence and metadata outputs without re-parsing the original JSON, JSONL, or FASTA files.

In addition to local workflow outputs, the evidence package includes `editorforge_external_online_sources_crosscheck`.csv. This table records the public structural, sequence, software, repository, and database sources used to ground the reproducibility statements in this manuscript. The table contains 14 external-source entries. These include the RCSB Protein Data Bank entry for PDB 4MH8, the public 4MH8 coordinate-download endpoint, the primary MMLV RT structural paper, the UniProt P03355 cross-reference, the ProteinMPNN publication and code repository, AlphaFold, ColabFold, and ESMFold literature/code sources, and general database context entries for RCSB PDB and UniProtKB [17, 18, 19, 20, 21, 4, 9, 10, 11]. A condensed version of this source crosscheck is provided in Table 4.

**Table 4:** Condensed external-source crosscheck for public structural, sequence, software, and database resources. The full table is provided as `editorforge_external_online_sources_crosscheck.csv`. The crosscheck records 14 public-source entries used to ground the MMLV RT target, the external inverse-folding engine, compatible structure-prediction/refolding routes, and public database context. This table is included to separate EditorForge-generated outputs from external resources used for target preparation, candidate generation, structural prediction, and reproducibility statements.

| Source ID | Category | Item | Use in EditorForge |
| --- | --- | --- | --- |
| EXT001 | Structural target | PDB 4MH8 / RCSB Protein Data Bank | Public structural source for the MMLV RT demonstration target |
| EXT002 | Structural target download | 4MH8 PDB coordinate file | Public coordinate source for reproducing target preparation |
| EXT003 | Primary literature | Das and Georgiadis 2004 MMLV RT structure paper | Primary literature reference for the MMLV RT structural target |
| EXT004 | Sequence annotation | UniProt P03355 | Public sequence/annotation cross-reference for the target protein family |
| EXT005 | Inverse-folding model | ProteinMPNN paper | Fixed-backbone sequence-generation method wrapped by EditorForge constraints |
| EXT006 | Inverse-folding code | ProteinMPNN GitHub repository | External code dependency for constrained sequence generation |
| EXT007 | Structure-prediction model | AlphaFold paper | Contextual structure-prediction reference for downstream refolding QC |
| EXT008 | Structure-prediction code | AlphaFold open-source code | Optional external predictor source for predicted PDB generation |
| EXT009 | Structure-prediction model | ColabFold paper | Compatible route for generating predicted structures consumed by RefoldQC |
| EXT010 | Structure-prediction code | ColabFold GitHub repository | External structure-prediction workflow compatible with RefoldQC handoff |
| EXT011 | Structure-prediction model | ESMFold paper | Alternative single-sequence structure-prediction source compatible with RefoldQC |
| EXT012 | Structure-prediction code | ESM / ESMFold GitHub repository | Optional source for predicted PDB generation if ESMFold outputs are supplied |
| EXT013 | Database context | RCSB PDB database | Public database context for parent structural data |
| EXT014 | Database context | UniProtKB database | Public sequence/annotation database context |

**Table 5:**
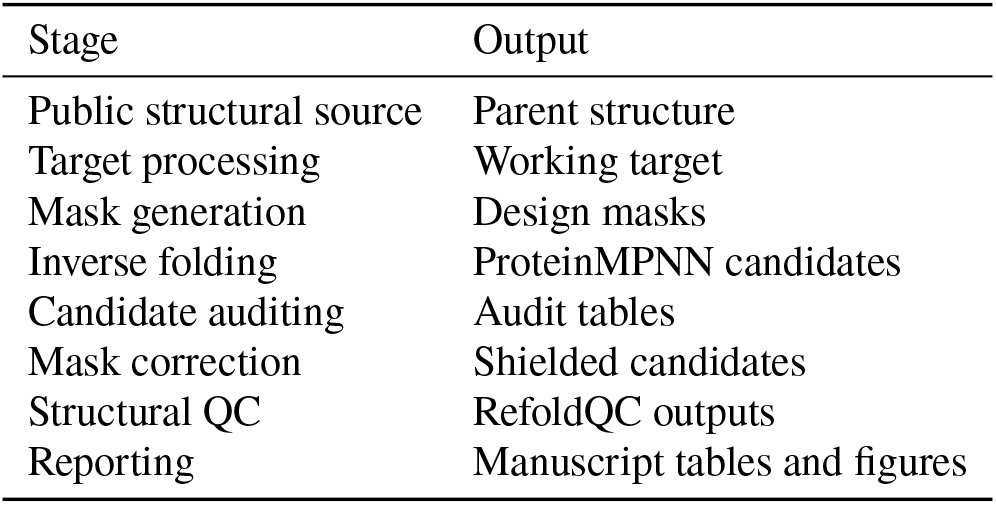
EditorForge reproducibility chain.

| Stage | Output |
| --- | --- |
| Public structural source | Parent structure |
| Target processing | Working target |
| Mask generation | Design masks |
| Inverse folding | ProteinMPNN candidates |
| Candidate auditing | Audit tables |
| Mask correction | Shielded candidates |
| Structural QC | RefoldQC outputs |
| Reporting | Manuscript tables and figures |

The external-source crosscheck supports three specific reproducibility claims. First, the parent structural target is publicly grounded as PDB 4MH8 is listed in the crosscheck table as the RCSB structural source for the MMLV RT demonstration, with the processed 453-residue EditorForge working target derived from that public structure[17, 16]. Second, the inverse-folding engine is externally grounded: ProteinMPNN is listed as the fixed-backbone sequence-generation method wrapped by EditorForge masks and audits. Third, the structure-prediction/refolding handoff is externally grounded: AlphaFold-style workflows, ColabFold, and ESMFold are listed as compatible external routes for producing predicted PDB files that can be consumed by Enhanced RefoldQC [9, 10, 11]. These entries do not imply that EditorForge authorship extends to those external tools; they document the public resources and dependencies around which EditorForge was built.

The crosscheck table also prevents overstatement because for example, UniProt P03355 is included as a public sequence/annotation cross-reference for the target protein family, but the 453-residue EditorForge working FASTA is not treated as the same as the UniProt sequence [20, 21]. Similarly, AlphaFold, ColabFold, and ESMFold are included as compatible structure-prediction sources, but a specific predictor is identified as the actual refolding engine only when it was used for the returned predicted structures. This distinction is important because the present study evaluates the EditorForge triage workflow, not the development of a new inverse-folding or structure-prediction model.

The repository also includes the frozen `Important Data` evidence folder, the final figure assets, the manuscript-ready tables, the source-file copies, the figure/table captions, and the scripts used to regenerate the figures and tables from the frozen local outputs. The intended reproducibility chain is therefore:

This chain defines the scope of the available data. It supports reproducible computational triage and source verification rather than experimental validity

All public-source identifiers, URLs, database names, and literature identifiers used for external grounding are `recorded_in_editorforge_external_online_sources_crosscheck`.csv. All local raw-output provenance and schema metadata are recorded in `RAW_DATA_FILE_MANIFEST.csv` and `RAW_DATA_SCHEMA_DICTIONARY.csv`. Candidate-level numerical results in the main manuscript should be taken from TABLE 2_FINAL_CANDIDATE_DASHBOARD.csv. The remaining combined tables are supplementary repository data products intended to make the raw outputs inspectable and reusable.

## 7 Acknowledgements

We acknowledge the St. Andrew’s College and The Scientific Research Collective in Aurora, Canada for providing the academic environment and oversight in which this project was developed. In particular, we are extremely grateful for the support of Joel Morrissey, the director of the collective, as well as Dina Baker for her invaluable guidance, oversight, and support throughout this research, which was instrumental in the development of this paper. We also thank the broader research and computational biology communities whose open structural databases, software tools, and documentation made this work possible.

## 7.1 Author contributions

A. Chen conceived the EditorForge workflow, designed the constraint-and-audit framework, implemented the computational pipeline, co-created scripts, analyzed the data and results, wrote the Results and Discussion sections, and guided project execution/coordination.

J. Siddiqui heavily assisted with project execution/coordination, contributed to core data synthesis, co-developed the experimental methodology, and helped draft the segments of the manuscript

W. Taucar oversaw group quality assurance, conducted data reviews, and helped manage the creation of the reference library.

L. Tiralongo co-created script designs, helped develop the primary data-analysis codebase, and executed simulations used to validate the framework.

M. Tkachenko greatly assisted with manuscript development, structured editing, and synthesis of technical sections to improve narrative cohesion.

A. Xu contributed to the quantitative data-analysis workflow and also greatly assisted with statistical processing and simulation-script editing.

S. Bawa provided editorial support during the results and discussion sections.

S. Guo assisted with LaTeX editing and manuscript formatting.

O. Pinska performed targeted literature review to contextualize the findings regarding competing methodologies

J. Rim assisted with data analysis of simulation results and LaTeX code creation

J. Shi assisted with synthesis of claims as well as creation of nonclaims sections

M. Wang conducted foundational literature review for relevant background research.

E. Zhao assisted with design and production of manuscript figures and visual frameworks and conducted bench-marking checks against existing methodologies.

## Notes

https://github.com/anthonycehn-eng/EditorForge-An-Active-Site-Aware-Framework-for-Inverse-Folding-Based-Protein-Redesign.git

### Competing Interest Statement

The authors have declared no competing interest.

## References

[1] Yue, K. and Dill, K. A. Inverse protein folding problem: designing polymer sequences. Proceedings of the National Academy of Sciences 89, 4163–4167 (1992). doi:10.1073/pnas.89.9.4163.

[2] Ingraham, J., Garg, V. K., Barzilay, R. and Jaakkola, T. Generative models for graph-based protein design. Advances in Neural Information Processing Systems 32 (2019).

[3] Hsu, C., Verkuil, R., Liu, J., Lin, Z., Hie, B., Sercu, T., Lerer, A. and Rives, A. Learning inverse folding from millions of predicted structures. Proceedings of the 39th International Conference on Machine Learning, PMLR 162, 8946–8970 (2022).

[4] Dauparas, J., Anishchenko, I., Bennett, N., Bai, H., Ragotte, R. J., Milles, L. F., Wicky, B. I. M., Courbet, A., de Haas, R. J., Bethel, N., Leung, P. J. Y., Huddy, T. F., Pellock, S., Tischer, D., Chan, F., Koepnick, B., Nguyen, H., Kang, A., Sankaran, B., Bera, A. K., King, N. P. and Baker, D. Robust deep learning-based protein sequence design using ProteinMPNN. Science 378, 49–56 (2022). doi:10.1126/science.add2187.

[5] Tang, X., Ye, X., Wu, F., Shao, D., Fang, Y., Chen, S., Xu, D. and Gerstein, M. BC-Design: A biochemistry-aware framework for highly accurate inverse protein folding. bioRxiv (2025). doi:10.1101/2024.10.28.620755.

[6] Kuhlman, B. and Bradley, P. Advances in protein structure prediction and design. Nature Reviews Molecular Cell Biology 20, 681–697 (2019). doi:10.1038/s41580-019-0163-x.

[7] Alford, R. F., Leaver-Fay, A., Jeliazkov, J. R., O’Meara, M. J., DiMaio, F. P., Park, H., Shapovalov, M. V., Renfrew, P. D., Mulligan, V. K., Kappel, K., Labonte, J. W., Pacella, M. S., Bonneau, R., Bradley, P., Dunbrack, R. L., Das, R., Baker, D., Kuhlman, B., Kortemme, T. and Gray, J. J. The Rosetta all-atom energy function for macromolecular modeling and design. Journal of Chemical Theory and Computation 13, 3031–3048 (2017). doi:10.1021/acs.jctc.7b00125.

[8] Anishchenko, I., Pellock, S. J., Chidyausiku, T. M., Ramelot, T. A., Ovchinnikov, S., Hao, J., Bafna, K., Norn, C., Kang, A., Bera, A. K., DiMaio, F., Carter, L., Chow, C. M., Montelione, G. T. and Baker, D. De novo protein design by deep network hallucination. Nature 600, 547–552 (2021). doi:10.1038/s41586-021-04184-w.

[9] Jumper, J., Evans, R., Pritzel, A., Green, T., Figurnov, M., Ronneberger, O., Tunyasuvunakool, K., Bates, R., Zidek, A., Potapenko, A., Bridgland, A., Meyer, C., Kohl, S. A. A., Ballard, A. J., Cowie, A., Romera-Paredes, B., Nikolov, S., Jain, R., Adler, J., Back, T., Petersen, S., Reiman, D., Clancy, E., Zielinski, M., Steinegger, M., Pacholska, M., Berghammer, T., Bodenstein, S., Silver, D., Vinyals, O., Senior, A. W., Kavukcuoglu, K., Kohli, P. and Hassabis, D. Highly accurate protein structure prediction with AlphaFold. Nature 596, 583–589 (2021). doi:10.1038/s41586-021-03819-2.

[10] Mirdita, M., Schütze, K., Moriwaki, Y., Heo, L., Ovchinnikov, S. and Steinegger, M. ColabFold: making protein folding accessible to all. Nature Methods 19, 679–682 (2022). doi:10.1038/s41592-022-01488-1.

[11] Lin, Z., Akin, H., Rao, R., Hie, B., Zhu, Z., Lu, W., Smetanin, N., dos Santos Costa, A., Fazel-Zarandi, M., Sercu, T., Candido, S. and Rives, A. Evolutionary-scale prediction of atomic-level protein structure with a language model. Science 379, 1123–1130 (2023). doi:10.1126/science.ade2574.

[12] Baek, M., DiMaio, F., Anishchenko, I., Dauparas, J., Ovchinnikov, S., Lee, G. R., Wang, J., Cong, Q., Kinch, L. N., Schaeffer, R. D., Millan, C., Park, H., Adams, C., Glassman, C. R., DeGiovanni, A., Pereira, J. H., Rodrigues, A. V., van Dijk, A. A., Ebrecht, A. C., Opperman, D. J., Sagmeister, T., Buhlheller, C., Pavkov-Keller, T., Rathinaswamy, M. K., Dalwadi, U., Yip, C. K., Burke, J. E., Garcia, K. C., Grishin, N. V., Adams, P. D., Read, R. J. and Baker, D. Accurate prediction of protein structures and interactions using a three-track neural network. Science 373, 871–876 (2021). doi:10.1126/science.abj8754.

[13] Baek, M., McHugh, R., Anishchenko, I., et al. Accurate prediction of protein–nucleic acid complexes using RoseTTAFoldNA. Nature Methods (2024). doi:10.1038/s41592-023-02086-5.

[14] Anzalone, A. V., Randolph, P. B., Davis, J. R., Sousa, A. A., Koblan, L. W., Levy, J. M., Chen, P. J., Wilson, C., Newby, G. A., Raguram, A. and Liu, D. R. Search-and-replace genome editing without double-strand breaks or donor DNA. Nature 576, 149–157 (2019). doi:10.1038/s41586-019-1711-4.

[15] Shuto, Y., et al. Structural basis for pegRNA-guided reverse transcription by prime editor. Nature (2024). doi:10.1038/s41586-024-07497-8.

[16] Das, D. and Georgiadis, M. M. The crystal structure of the monomeric reverse transcriptase from Moloney murine leukemia virus. Structure 12, 819–829 (2004). doi:10.1016/S0969-2126(04)00115-7.

[17] RCSB Protein Data Bank. PDB ID 4MH8: The crystal structure of the monomeric reverse transcriptase from Moloney murine leukemia virus. RCSB PDB (accessed June 2026). doi:10.2210/pdb4MH8/pdb.

[18] Berman, H. M., Westbrook, J., Feng, Z., Gilliland, G., Bhat, T. N., Weissig, H., Shindyalov, I. N. and Bourne, P. E. The Protein Data Bank. Nucleic Acids Research 28, 235–242 (2000). doi:10.1093/nar/28.1.235.

[19] Burley, S. K., Berman, H. M., Bhikadiya, C., et al. RCSB Protein Data Bank: biological macromolecular structures enabling research and education in fundamental biology, biomedicine, biotechnology and energy. Nucleic Acids Research 47, D464–D474 (2019). doi:10.1093/nar/gky1004.

[20] The UniProt Consortium. UniProt: the Universal Protein Knowledgebase in 2025. Nucleic Acids Research 53, D609–D617 (2025). doi:10.1093/nar/gkae1010.

[21] The UniProt Consortium. UniProtKB entry P03355: Gag-Pol polyprotein, Moloney murine leukemia virus isolate Shinnick. UniProtKB (accessed June 2026).

[22] CÔté, M. L. and Roth, M. J. Murine leukemia virus reverse transcriptase: structural comparison with HIV-1 reverse transcriptase. Virus Research 134, 186–202 (2008). doi:10.1016/j.virusres.2008.01.001.

[23] Sarafianos, S. G., Marchand, B., Das, K., Himmel, D. M., Parniak, M. A., Hughes, S. H. and Arnold, E. Structure and function of HIV-1 reverse transcriptase: molecular mechanisms of polymerization and inhibition. Journal of Molecular Biology 385, 693–713 (2009). doi:10.1016/j.jmb.2008.10.071.

[24] Kabsch, W. A solution for the best rotation to relate two sets of vectors. Acta Crystallographica Section A 32, 922–923 (1976). doi:10.1107/S0567739476001873.

[25] Kabsch, W. A discussion of the solution for the best rotation to relate two sets of vectors. Acta Crystallographica Section A 34, 827–828 (1978). doi:10.1107/S0567739478001680.

[26] Umeyama, S. Least-squares estimation of transformation parameters between two point patterns. IEEE Transactions on Pattern Analysis and Machine Intelligence 13, 376–380 (1991). doi:10.1109/34.88573.

[27] Lawrence, J., Bernal, J. and Witzgall, C. A purely algebraic justification of the Kabsch-Umeyama algorithm. Journal of Research of the National Institute of Standards and Technology 124, 124028 (2019). doi:10.6028/jres.124.028.

[28] Wilkinson, M. D., Dumontier, M., Aalbersberg, I. J., et al. The FAIR Guiding Principles for scientific data management and stewardship. Scientific Data 3, 160018 (2016). doi:10.1038/sdata.2016.18.

[29] Barker, M., Chue Hong, N. P., Katz, D. S., Lamprecht, A.-L., Martinez-Ortiz, C., Psomopoulos, F., Harrow, J., Castro, L. J., Gruenpeter, M., Martinez, P. A. and Honeyman, T. Introducing the FAIR Principles for research software. Scientific Data 9, 622 (2022). doi:10.1038/s41597-022-01710-x.

[30] Sandve, G. K., Nekrutenko, A., Taylor, J. and Hovig, E. Ten simple rules for reproducible computational research. PLOS Computational Biology 9, e1003285 (2013). doi:10.1371/journal.pcbi.1003285.

